# Conversational production and comprehension: fMRI-evidence reminiscent of the classic Broca-Wernicke model

**DOI:** 10.1101/2023.07.05.547796

**Authors:** Caroline Arvidsson, Ekaterina Torubarova, André Pereira, Julia Uddén

**Affiliations:** Department of Linguistics, Stockholm University, Universitetsvägen 10 C, 114 18, Stockholm Sweden; Division of Speech, Music, and Hearing, KTH Royal Institute of Technology, Lindstedtsvägen 24, 114 28, Stockholm, Sweden; Department of Psychology, Stockholm University, Albanovägen 12, 114 19, Stockholm, Sweden

**Keywords:** Interaction, contextual language processing, LIFG, LMTG, functional asymmetry

## Abstract

A key question in neurolinguistics is whether language production and comprehension share neural infrastructure, but this question has not been addressed in the context of actual conversation. We utilized a public fMRI dataset where participants (N=24) engaged in unscripted conversations with a confederate outside the scanner via an audio-video link. We provide evidence indicating that production and comprehension, in a conversational setting, diverge with respect to how they modulate the recruitment of regions in the left-lateralized perisylvian language network. Activity in the left inferior frontal gyrus was stronger in production than in comprehension. Compared to production, comprehension showed stronger recruitment of the left anterior middle temporal gyrus and superior temporal sulcus, but this was not the case for the posterior aspect of these loci. Although our results are reminiscent of the classic Broca-Wernicke model, the anterior temporal activation is a notable difference from that model. This is one of the findings which may be a consequence of the conversational setting, another being that conversational production activated what we interpret as higher-level socio-pragmatic processes. In conclusion, we present evidence supporting that the above-mentioned frontal vs temporal regions in the language network are functionally segregated during conversation.

---

Conversation is integral to the everyday experience of almost every human. It is thus not surprising that language processing, as occurring during conversation, is the dominant explanandum in psycho- and neurolinguistics. Notwithstanding, isolation paradigms, where participants produce or listen to linguistic signals in a non-interactive setting, are standard in behavioral and in particular neuroimaging experiments. Conversation entails flexibly shifting between speaker and listener roles (Levinson and Torreira, 2015; Sacks et al., 1978), while simultaneously considering linguistic, social, and other contextual factors to encode and decode meaning (Austin, 1973; Grice, 1975). As all of that is missing in the isolation paradigm, one can question the validity of typical psycho-or neurolinguistic experiments in relation to the explanandum. The goal of this fMRI study was to address this issue by investigating the underlying processes of speech production and comprehension during actual conversation.

A long-standing but still ongoing debate is to what extent production and comprehension diverge with respect to the recruitment of regions in the perisylvian language network (Giglio et al., 2022b; Hu et al., 2022; Matchin et al., 2022; Matchin and Hickok, 2020; Rutten, 2022; Tremblay and Dick, 2016; Giglio et al., 2022a). The perisylvian language network (Caplan, 1987) is a left hemisphere-dominant network of cortical areas including the left inferior frontal gyrus (LIFG), the left middle/superior temporal gyri (LMTG/STG) and the posteroinferior parietal cortex. It is known to be crucial to higher-level linguistic processing, e.g., syntactic and semantic processing on the sentence level (Fedorenko et al., 2011; Hickok, 2022; Mollica et al., 2020; Stockert et al., 2023; Tyler et al., 2010; Vlooswijk et al., 2010; Malik-Moraleda et al., 2022). According to the classic model of the neurobiology of language, originating from the pioneering work of Carl Wernicke in the late 19th century, ‘Broca’s area’ (the posterior LIFG, following Paul Broca) was described as a motor speech center, and ‘Wernicke’s area’ (the posterior LSTG) a sensory speech center (Geschwind, 1970; Rutten, 2022; Tremblay and Dick, 2016; Wernicke, 1885/1977). At variance with the classic model were later lesion and neuroimaging studies, providing compelling evidence that frontal and temporal regions both subserve aspects of production and comprehension (Caramazza and Zurif, 1976; Fridriksson et al., 2015; Segaert et al., 2012; Hagoort, 2016). However, questions on the relative contribution of language-supporting regions to the production and comprehension systems (Pickering and Garrod, 2007) remain unresolved.

Hu et al. (2022) argue for a model in which production and comprehension rely on the same knowledge representations (see also Chomsky, 2014; Pickering and Garrod, 2004) and, by extension, the same neural structures. In an experimentally controlled fMRI study, these authors found no evidence of brain regions within or outside the language network that selectively supported processing in one of the systems (production or comprehension) but not the other. Hu et al. (2022) also found that all language regions (localized by contrasting reading sentences vs lists of nonwords) were more engaged during production than during comprehension. However, the production and comprehension processes tapped by the Hu et al. (2022) tasks likely differ from conversational language processes. For example, their comprehension tasks were likely less demanding than conversational comprehension, since they involved reading or listening to context-independent sentences containing single clauses (e.g., ‘the girl is smelling a flower’) and did not require the listener to integrate both previous linguistic and other contextual information to understand the message.

Conversely, Matchin and Hickok (2020) argue for a functional division of inferior frontal and temporal regions. According to their model, the posterior LIFG supports processes important to production specifically, namely the transformation of abstract morphosyntactic representations into linear sequences of morphemes, while the left posterior middle temporal gyrus (LpMTG) connects conceptual-semantic systems in the temporal and inferior parietal lobes during both production and comprehension. Moreover, it is possible that posterior aspects of the temporal cortex play a particularly crucial role in comprehension; in a lesion-to-symptom mapping study, Matchin et al. (2022) found that damage to the posterior middle temporal gyrus was mainly associated with syntactic comprehension deficits, while damage to the posterior inferior frontal gyrus was linked to expressive agrammatism.

A functional asymmetry of inferior frontal and temporal regions was indeed observed in a recent study conducted by Giglio et al. (2022b). In their study, participants listened to and produced word sequences of fixed lengths. Following the classical paradigm in Pallier et al. (2011), the words comprised phrases of varying sizes. Giglio et al. (2022b) found an effect of constituent size in frontal and temporal regions for both production and comprehension. However, when contrasting production and comprehension (all constituent sizes), activation of inferior frontal regions was stronger for production, while activation of middle temporal regions was stronger for comprehension. Results from an ROI analysis in Giglio et al. (2022b) also indicated that production entails stronger LIFG activity than comprehension, while the opposite was the case in the LMTG (i.e., more activity for comprehension than production). However, Giglio et al. (2022b) used large ROIs (’functional masks’) and their LIFG mask also covered regions outside of the LIFG. This use of large LIFG masks when comparing production with comprehension is problematic because areas adjacent to the IFGL (e.g., anterior insula and the MFGL/inferior frontal sulcus) are part of the multiple demand (MD) network (Camilleri et al., 2018; Duncan, 2013; MacGregor et al., 2022; Stiers et al., 2010; Wehbe et al., 2021). The MD network plays a crucial role in a set of domain-general processes often denoted by the umbrella term cognitive control or executive functions (e.g., working memory, inhibition, cognitive flexibility) (Miller and Cohen, 2001) which are recruited while speaking but do not primarily support linguistic processing (Diachek et al., 2020). In other words, the production activation in Giglio et al. (2022b) may simply reflect an increase in domain-general cognitive demand (Hu et al., 2022). Moreover, whether there is a division of labor in the perisylvian language network at all has yet to be tested within the context of actual conversation.

Presumably, all details of the full syntactic representation need to be included in the conversational production (Bock, 1982; Garrett, 1988). Conversational comprehension, on the other hand, relies on prediction-related processes that undermine the need for a full syntactic parse (Christianson et al., 2001; Ferreira, 2003; Ferreira et al., 2001; Ferreira and Patson, 2007; Townsend and Bever, 2001). The propensity to employ such mechanisms that increase processing speed should increase in conversational comprehension because of the demands of timing in turn-taking (Levinson and Torreira, 2015; Heldner and Edlund, 2010). Under this explanation, one would expect an even stronger asymmetry than previously observed (Giglio et al., 2022b) when comparing the neural responses of conversational production with such responses of conversational comprehension.

Conversation also involves socio-pragmatic processes (Austin, 1973; Grice, 1975; Wilson and Sperber, 2002) above and beyond word and utterance/sentence-level phonology, syntax, and semantics. These processes have however not previously been localized for production and comprehension in a conversational context. Rauchbauer et al. (2019) explored the neural correlates of human-human and human-robot interaction by modeling entire conversations (i.e., without modeling production and comprehension as separate regressors). In their study, conversation generated activation of the perisylvian language network, but also the dorsal frontal cortex, the temporo-parietal junction, and the occipital cortex, suggesting that interactive language use involves extra-linguistic networks related to the online tracking of one’s interlocutor’s mental state and facial gestures. Hogenhuis and Hortensius (2022) used the same dataset as in Rauchbauer et al. (2019) and modeled conversational production and comprehension separately. However, the purpose of the Hogenhuis and Hortensius (2022) study was to investigate how the tasks differed in human-human vs human-robot interaction, and although the visualization of their whole-brain analysis suggests that there are significant differences between production and comprehension in human-human interaction, they do not report on this explicitly in their study. We differ from Hogenhuis and Hortensius (2022) by (1) asking questions about production vs comprehension during human-human interaction only, (2) performing an ROI analysis similar to the ones that have been used in studies that address our research question (Giglio et al., 2022b; Hu et al., 2022), and (3) interpreting these results in terms of existing models on production vs comprehension.

In this fMRI study, we revisit the classic question of the relative contribution of frontal and temporal regions in the left-lateralized perisylvian language network, now in the context of actual conversation. In modern research on the neurobiology of language, divisions are made between the anterior and posterior aspects of both of these nodes (e.g., Matchin and Hickok, 2020; Matchin et al., 2022; Hu et al., 2022). As we were interested in these differences, we chose an ROI-approach (also following Giglio et al., 2022b), in addition to a whole-brain analysis.

## Materials and Methods

### Data

Raw MRI images and TextGrid-formatted orthographic transcriptions were retrieved from a publicly available data set provided by Rauchbauer et al. (2020). The MRI data and transcriptions were retrieved from OpenNeuro and Otrolang (https://hal.archives-ouvertes.fr/hal-02612820/, https://www.ortolang.fr/market/ corpora/convers/v2). The 25 participants in the corpus reported normal or corrected-to-normal vision and had no prior history of psychiatric or neurological conditions. One of these participants was excluded from the present study because of excessive head movement (movement *>* 4mm). Included in the main analysis were 24 participants (18 female, 6 male, M age = 28.8, SD = 12).

In the Rauchbauer et al. (2020) corpus, participants held conversations in their L1 (French) with a confederate in the control room. The confederate was either an experimenter or a robot (controlled by the experimenter through a Wizard of Oz paradigm), but in the present study, we were only interested in human-human conversation. We, therefore, modeled the images acquired during the human-robot conversations in the same way (using the same categories, see below) but separately from the images acquired during the human-human conversations. In the present study, only data from human-human conversations (12 min/participant in total) were used in the first-level contrasts and the second-level analysis. Henceforth, all the mentioned events refer to those of the human-human conversations.

Interlocutors were connected via bidirectional audio (using active noise-cancellation), and unidirectional video transmission (the participant saw the confederate’s face on a video monitor, but not vice versa). To provide a framework for naturalistic conversation, participants were told that they would discuss images from an advertising campaign with another participant. Rauchbauer et al. (2020) reported that all participants confirmed that they believed the cover story after participation. There were four runs per participant, each consisting of six blocks with the following block structure: 8-sec presentation of the image of the fruit, 4-sec fixation cross, 1 min conversation with the confederate, 4-sec fixation cross.

Rauchbauer et al. (2020) collected MRI data with a 3T Siemens Prisma and a 20-channel head coil. Functional images were acquired using an EPI sequence with the following parameters: echo time (TE): 30 ms, repetition time (TR): 1205 ms, matrix size: 84 × 84, field of view (FOV): 210 mm × 210 mm, voxel size (VS): 2.5 × 2.5 × 2.5 mm^3^, 54 slices co-planar to the anterior/posterior commissure plane (axial), flip angle: 65°. Functional images were acquired with multiband acquisition factor 3. Parameters for the acquisition of structural images were: TE: 0.002 ms, TR: 2.4 ms, FOV: 204.8 × 256 × 256 mm, VS: 0.8 × 0.8 × 0.8 mm^3^, 320 slices (sagittal).

Rauchbauer et al. (2020) automatically segmented audio files of speech from individual speakers into inter-pausal units (blocks of speech surrounded by silences *≥* 200 ms) that were visually inspected and manually transcribed. In the present study, we extracted onsets and offsets of three events: production (when the participant spoke), comprehension (when the confederate spoke), and silence (when both interlocutors were silent), from the transcribed data in Rauchbauer et al. (2020). This extraction was performed using a Python script (https://github.com/ carolinearvidsson/RobotfMRI). To avoid extremely short events in the analysis, utterances were merged into a single utterance if they were surrounded by silences *<* 300 ms within the same speaker. Utterances shorter than 300 ms were removed.

### fMRI Preprocessing

We performed rigid body transformation using 6 parameters (translations and rotations). Head movements in coordinates x, y, and z were inspected independently. As previously mentioned, one participant had *>* 4 mm head movement and was therefore excluded from the following analyses. Functional images were coregistered to an anatomical image (T1) and normalized to a standard MNI space with affine regularization. Normalization included a resampling of the voxels to 2 x 2 x 2 mm with a 4th-degree B-spline interpolation. White and grey matter segmentation and bias correction were conducted during the normalization step. Finally, functional images were spatially smoothed using a 3D isotropic 5 mm full-with-at-half-maximum Gaussian kernel. A temporal high-pass filter (cycle cut-off at 128 sec) was used to account for low-frequency effects.

### Whole-brain analysis

For the first-level single-subject analysis, production, comprehension, and silences were modeled as three separate regressors. Images acquired during the presentation of the fixation cross (fixation) and the presentation of the advertising image (advertisement) were modeled as two separate regressors. Head movements were modeled as six motion parameters. The events were convolved with a canonical hemodynamic response function. The three regressors used in the contrasts were production, comprehension, and fixation. Production and comprehension were contrasted against fixation and each other: production *>* fixation, comprehension *>* fixation, production *>* comprehension, comprehension *>* production. The number and duration of the production and comprehension events are available in Table 1. The second level analysis was conducted with one-sample t-tests on the contrast images defined at the first level. A cluster-forming threshold of *p_uncorrected_* was set to .001 (no extent-level threshold, *k* = 0). Family-wise error (FWE), as implemented in SPM12, was used as the multiple comparison correction method (cluster and peak level). Only clusters with *p_F W E_ <* .05 at cluster level were reported in the current investigation. The test statistic of each cluster’s highest peak (voxel) is also reported. No additional voxels were reported, even if they were significant at *p_F W E_ <* 0.05 at the voxel level. Cluster labeling was performed using the Automated anatomical labeling atlas toolbox for SPM (Rolls et al., 2020).

**Table 1.** The number of production and comprehension events. The mean, SD, and range of the event durations are given in seconds. Events shorter than 0.3 secs were removed from the analysis.

|  | Production | Comprehension |
| --- | --- | --- |
| N | 3447 | 3538 |
| Mean | 1.67 | 1.96 |
| SD | 1.30 | 1.56 |
| Range | 0.3-9.96 | 0.3-10.43 |

### ROI analysis

Anatomical ROIs (see Figure 3) were extracted from the Harvard-Oxford cortical structural atlas (HO atlas) (downloaded from Neurovault in April 2023: https://identifiers.org/neurovault.image:1702). The anatomical ROIs enabled us to focus on the same areas as in Giglio et al. (2022b), who also investigated the division of labor of the LIFG and the LMTG during production and comprehension, without having the masks extending into regions outside of the inferior frontal and middle temporal gyri and adjacent sulci. Moreover, the LpMTG/STG have traditionally been the main focus in the debate regarding the distribution of labor between frontal and temporal regions (Matchin et al., 2022; Rutten, 2022), but the whole-brain results from Giglio et al. (2022b) suggested rather that activation of more anterior and middle parts of the LMTG/STG are stronger for comprehension than production. There are also empirical reasons to assume that there is a functional division of the LIFG subregions, e.g., that anterior aspects facilitate semantic processes, while more posterior aspects facilitate phonological processes (for a review, see Bookheimer, 2002). Thus, we divided the LIFG and LMTG into the following subregions: the LIFG pars orbitalis (LIFGOrb), pars triangularis (LIFGTri), and pars opercularis (LIFGOper) and the anterior LMTG (LaMTG, including the left anterior superior temporal sulcus; LaSTS), and the LpMTG (including the LpSTS). The LIFGOrb was not labeled in the HO atlas. To extract only the LIFGOrb, the orbitofrontal cortex in the HO atlas was masked with the LIFGOrb as defined in the Automated anatomical labeling atlas (AAL3v1, downloaded from https://www.oxcns.org/aal3.html). The mean beta weights per ROI and participant were calculated in SPM for contrasts production *>* fixation and comprehension *>* fixation.

With the extracted beta values from the anatomical ROIs described above, we ran a linear mixed effects model to investigate the interaction effect of LOBE (frontal, temporal) and SYSTEM (production, comprehension), with by-participant random intercepts. The linear mixed models were conducted using the lme4 package (Vazquez et al., 2010) in R (R Core Team, 2021), with an alpha level of *α* = 0.05. P-values were retrieved using R package afex (Singmann et al., 2018). One sample t-tests were conducted to investigate whether the mean beta weights differed from zero. Paired sample t-tests were conducted to compare ROI recruitment in production with comprehension. The p-values of the t-tests were adjusted for multiple comparisons (Bonferroni correction for five comparisons, i.e., one comparison per ROI).

## Results

### Whole-Brain Analysis

We investigated the main effect of production and comprehension against the baseline (looking at a fixation cross). The contrasts production *>* fixation generated four clusters: a cluster in the left hemisphere, spanning over the superior and middle temporal gyri (STG/MTG), the left pre/postcentral gyrus (PreCG/PoCG) all the way to the LIFG (including the pars orbitalis, pars triangularis and pars opercularis); a right-hemispheric cluster, covering the right STG, the right PreCG, and PoCG; a bilateral cluster in the supplementary motor area (SMA); and finally a bilateral cluster spanning from the right MTG/STG to the left occipital gyrus. The contrast comprehension *>* fixation generated five clusters: two clusters (one left-lateralized, one right-lateralized) spanned from the anterior aspects of the middle and superior temporal lobes to the insula; two clusters respectively covering the right and left occipital gyri; and one cluster in the LIFG pars triangularis (see Fig. 1, panel A and Table 2).

**Fig. 1:**
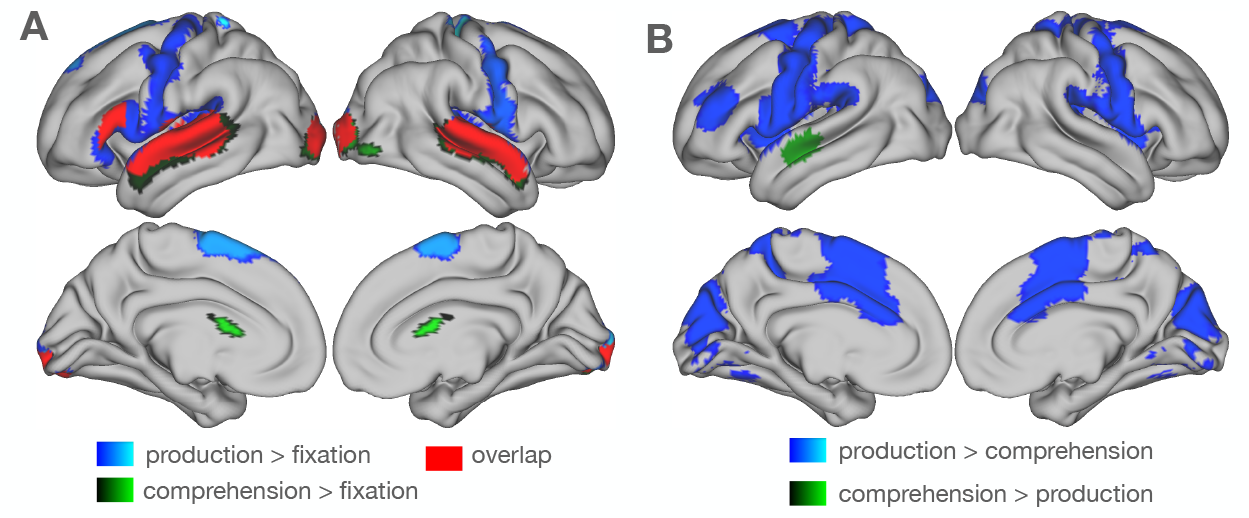
A: Whole-brain results for the main effect of systems (production, comprehension). Blue: areas active in production. Green: areas active in comprehension. Red: production and comprehension activation overlap. B: Whole-brain results from contrasting systems against each other. Blue: areas more active in production than comprehension. Green: areas more active in comprehension than production. The figure shows clusters with a cluster-forming threshold of *p_uncorrected_* = .001 (no extent-level threshold, k = 0). Only clusters with a *p_F W E_ <* .05 are reported.

**Table 2.**
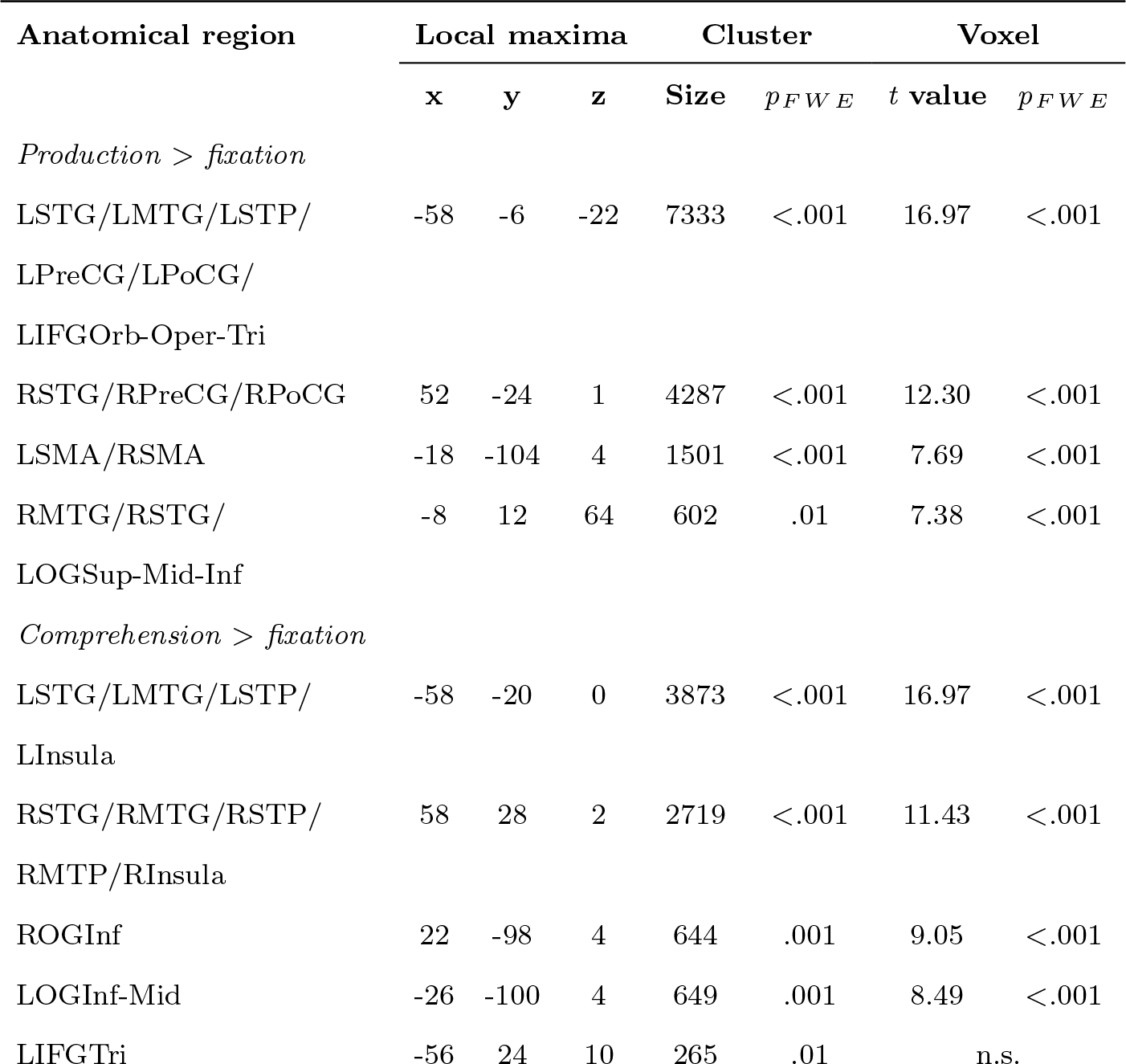
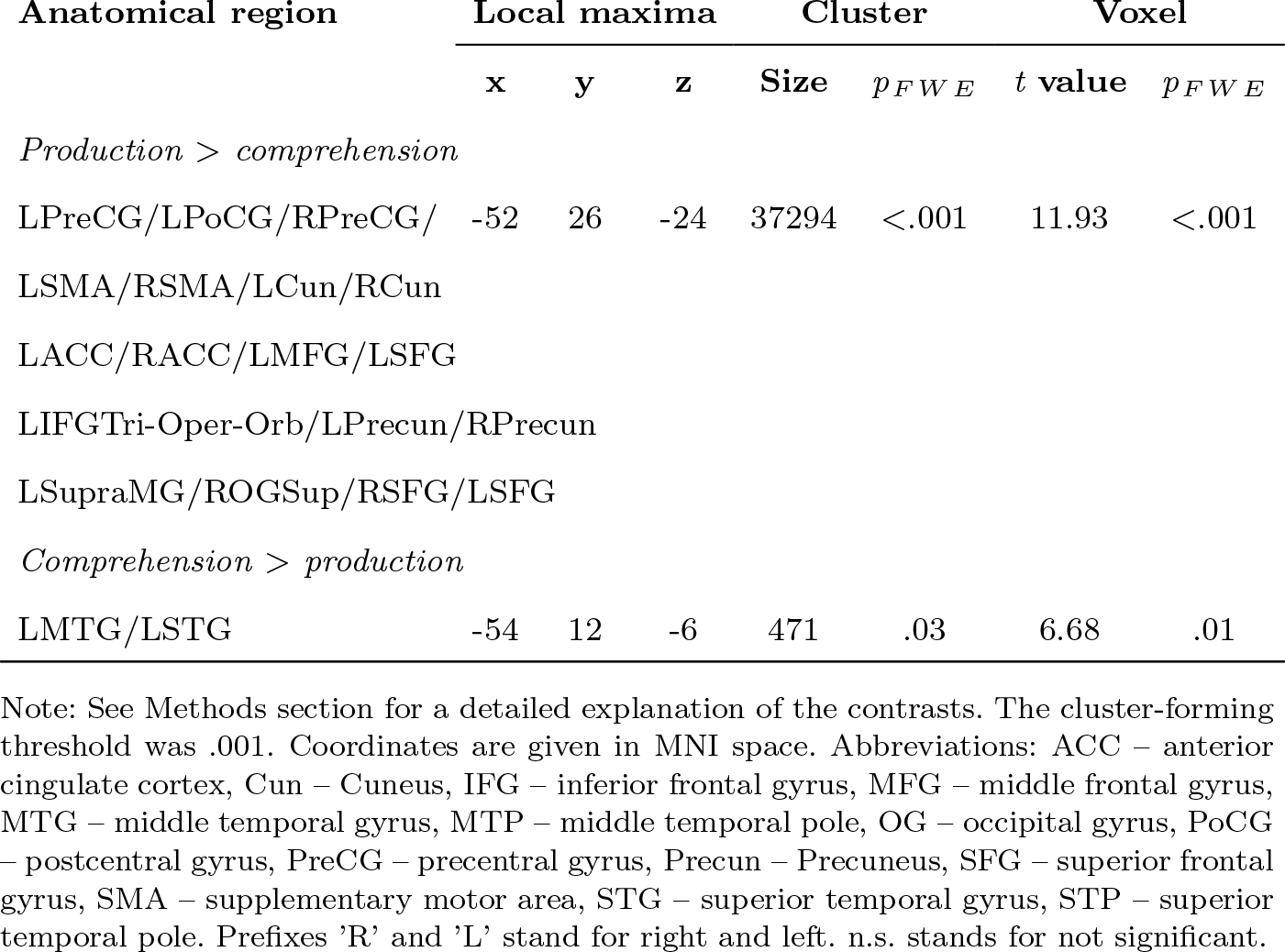
Activations for the contrasts production *>* fixation, comprehension *>* fixation, production *>* comprehension, comprehension *>* production. The number of voxels is given for each cluster, together with the MNI coordinates and t-value of the cluster’s maximum peak (local maxima).

| Anatomical region | Local maxima |  |  | Cluster |  | Voxel |  |
| --- | --- | --- | --- | --- | --- | --- | --- |
| | x | y | z | Size | $p_{FWE}$ | t value | $p_{FWE}$ |
| *Production > fixation* |  |  |  |  |  |  |  |
| LSTG/LMTG/LSTP/<br>LPreCG/LPoCG/<br>LIFGOrb-Oper-Tri | -58 | -6 | -22 | 7333 | <.001 | 16.97 | <.001 |
| RSTG/RPreCG/RPoCG | 52 | -24 | 1 | 4287 | <.001 | 12.30 | <.001 |
| LSMA/RSMA | -18 | -104 | 4 | 1501 | <.001 | 7.69 | <.001 |
| RMTG/RSTG/<br>LOGSup-Mid-Inf | -8 | 12 | 64 | 602 | .01 | 7.38 | <.001 |
| *Comprehension > fixation* |  |  |  |  |  |  |  |
| LSTG/LMTG/LSTP/<br>LInsula | -58 | -20 | 0 | 3873 | <.001 | 16.97 | <.001 |
| RSTG/RMTG/RSTP/<br>RMTP/RInsula | 58 | 28 | 2 | 2719 | <.001 | 11.43 | <.001 |
| ROGInf | 22 | -98 | 4 | 644 | .001 | 9.05 | <.001 |
| LOGInf-Mid | -26 | -100 | 4 | 649 | .001 | 8.49 | <.001 |
| LIFGTri | -56 | 24 | 10 | 265 | .01 | n.s. |  |
| | x | y | z | Size | $p_{FWE}$ | $t$ value | $p_{FWE}$ |
| *Production > comprehension* |  |  |  |  |  |  |  |
| LPreCG/LPoCG/RPreCG/<br>LSMA/RSMA/LCun/RCun<br>LACC/RACC/LMFG/LSFG<br>LIFGTri-Oper-Orb/LPrecun/RPrecun<br>LSupraMG/ROGSup/RSFG/LSFG | -52 | 26 | -24 | 37294 | <.001 | 11.93 | <.001 |
| *Comprehension > production* |  |  |  |  |  |  |  |
| LMTG/LSTG | -54 | 12 | -6 | 471 | .03 | 6.68 | .01 |

Note: See Methods section for a detailed explanation of the contrasts. The cluster-forming threshold was .001. Coordinates are given in MNI space. Abbreviations: ACC – anterior cingulate cortex, Cun – Cuneus, IFG – inferior frontal gyrus, MFG – middle frontal gyrus, MTG – middle temporal gyrus, MTP – middle temporal pole, OG – occipital gyrus, PoCG – postcentral gyrus, PreCG – precentral gyrus, Precun – Precuneus, SFG – superior frontal gyrus, SMA – supplementary motor area, STG – superior temporal gyrus, STP – superior temporal pole. Prefixes ‘R’ and ‘L’ stand for right and left. n.s. stands for not significant.

We also investigated areas more strongly recruited in production than comprehension and vice versa. The contrast production *>* comprehension generated a very large bilateral cluster, covering motor regions, the left inferior/left middle/bilateral superior frontal gyrus, the bilateral precuneus, the medial prefrontal cortex (mPFC), and the occipital lobe. Moreover, a rather anterior portion of the LMTG/STG was more activated for comprehension than production (see Fig. 1, panel B).

### ROI Analysis

All 10 distributions of mean beta values (one distribution for production and one for comprehension in each ROI) individually passed a Shapiro-Wilks normality test (sig. values were larger than 0.9). The mixed effects model with LOBE and SYSTEM as factors and participants as random effects showed that the effect of LOBE was significant, so that temporal lobe activation was stronger than frontal lobe activation (*β* = 2.17, SE = 0.29, *t*(213) = 7.61, p *<* .001). Activation overall was stronger in production than comprehension (*β* = 1.37, SE = 0.26, *t*(213) = 5.36, p *<* .001). We report these results for completeness, although we set up the model mainly to test the interaction. There was a significant interaction between LOBE and SYSTEM (see Fig. 2), showing opposite signs of LOBE activity differences (LIFG vs LMTG/STS), for production and comprehension (*β* = −2.55, SE = 0.40, *t*(213) = −6.33, p *<* .001). The mean beta values in production and comprehension differed significantly from each other in all regions except for the LpMTG (*t*(23) = −1.93, p = .33). Production activation was larger than comprehension activation in the LIFG pars orbitalis (*t*(23) = 3.63, p *<* 0.01), the LIFG pars triangularis (*t*(23) = 2.83, p *<* 0.05), and the LIFG pars opercularis (*t*(23) = 3.63, p *<* 0.01), while comprehension activation was larger than production activation in the LaMTG (*t*(23) = −3.51, p *<* 0.01). See Fig. 3.

**Fig. 2:**
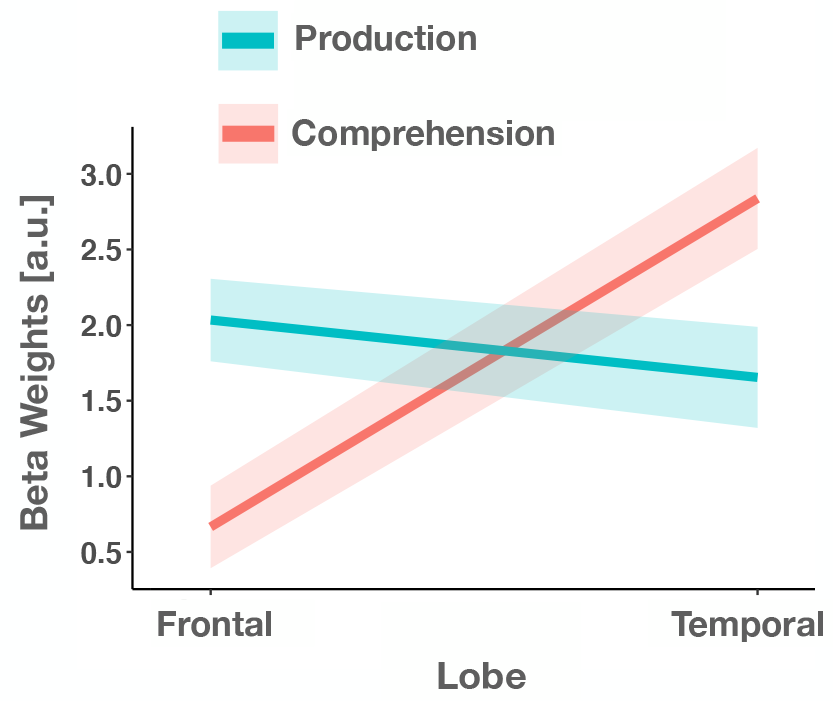
Interaction of LOBE (frontal, temporal) and SYSTEM (production, comprehension). Lines show differences between frontal and temporal activation in production (blue line) and comprehension (red line). Line ribbons show the standard error of the mean. Regions in frontal: LIFG pars orbitalis, triangularis, and opercularis. Regions in temporal: anterior and posterior LMTG/STS.

**Fig. 3:**
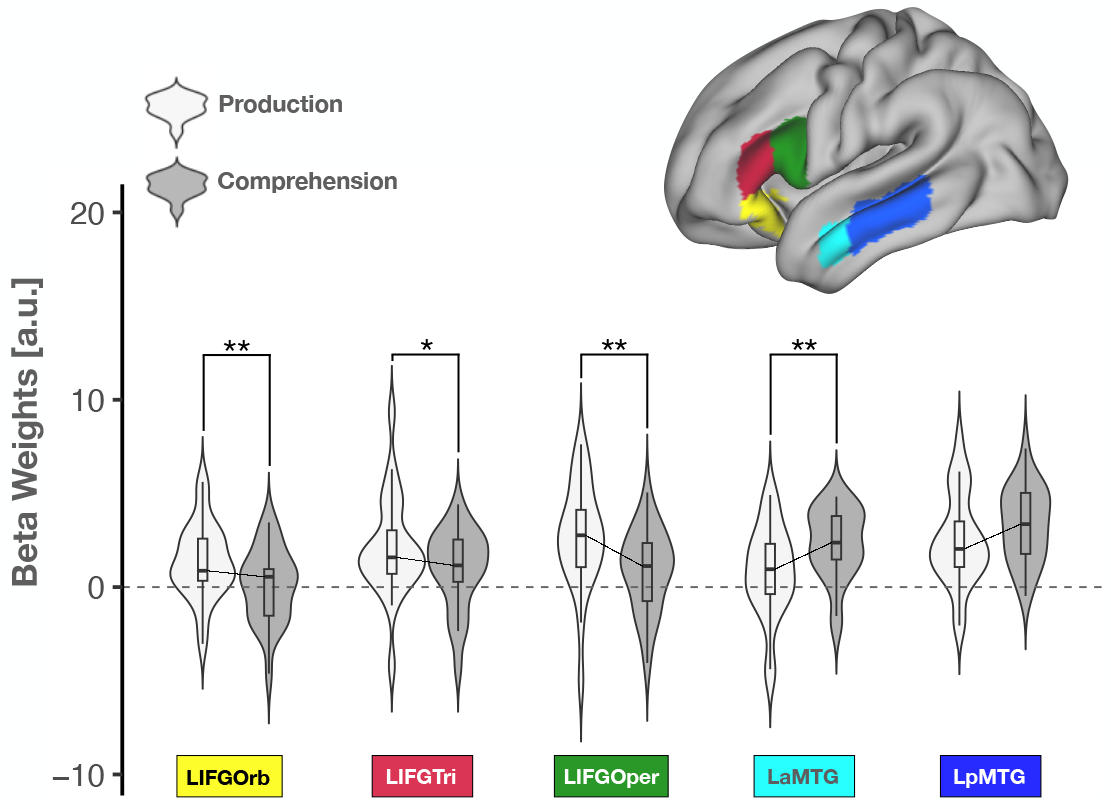
ROI-wise differences in BOLD response between conversational production and conversational comprehension. Each violin shows the density distribution of the participants’ mean parameter estimates from the contrasts production/comprehension vs baseline (white: production, gray: comprehension) and ROI. Yellow: left inferior frontal gyrus (LIFG) pars orbitalis, magenta: LIFG pars triangularis, green: LIFG pars opercularis, azure: left anterior middle temporal gyrus, extending into the left anterior superior temporal sulcus (LaMTG/STS), dark blue: left posterior LpMTG/STS. The lines reaching from one violin to another show the difference in means across tasks within each ROI. Star notation: * = p *<* .05; ** = p *<* .01. P-values were corrected for multiple comparisons.

In production, mean beta values were significantly different from zero in all ROIs, except for the LaMTG (LIFGorb: *t*(23) = 3.29, p = .02; LIFGtri: *t*(23) = 3.81, p *<* .01; LIFGoper: *t*(23) = 4.63, p *<* .001; LaMTG: *t*(23) = 1.66, p *<* 0.05; LpMTG: *t*(23) = 4.96, p *<* .001). For comprehension, mean beta values were significantly different from zero in the LaMTG, and the LpMTG, but not the LIFG pars orbitalis, the LIFG pars triangularis or the LIFG pars opercularis (LIFGorb: *t*(23) = 0.20, p *<* .05; LIFGtri: *t*(23) = 2.38, p *<* .05; LIFGoper: *t*(23) = 2.04, p *<* .05; LaMTG: *t*(23) = 2.04, p *<* .001; LpMTG: *t*(23) = 8.19, p *<* .001).

## Discussion

Using a neuroimaging paradigm where participants engaged in unscripted conversations, we have provided evidence that conversational production and conversational comprehension diverge with respect to the recruitment of the left-lateralized perisylvian language network. Our whole-brain results show that albeit both inferior frontal and superior/middle temporal regions subserve conversational production and comprehension, the recruitment of left inferior frontal gyrus (LIFG) regions was stronger in production than in comprehension, while the recruitment of the anterior aspects of the temporal lobe (the left anterior superior temporal sulcus, LaSTS; the left anterior superior temporal gyrus, LaSTG; aspects of the left anterior middle temporal gyrus, LaMTG) was stronger in comprehension than in production. The asymmetric recruitment during conversational production and conversational comprehension found further support in our ROI analysis, showing significantly greater activation of the LIFG and its individual subregions (pars orbitalis, pars triangularis, and pars opercularis) during production than comprehension. In turn, the ROI analysis showed that the LaMTG/STS plays a unique role for comprehension processes. The ROI analysis did not show a statistically significant difference between production and comprehension in the level of recruitment of the more posterior aspects of the left middle temporal gyrus/sulcus (LpMTG/STS). While these results will certainly remind the reader of the classic model (Geschwind, 1970; Rutten, 2022; Tremblay and Dick, 2016; Wernicke, 1885/1977), this more anterior location of the temporal activation for conversational comprehension is a crucial difference relative to that model (further discussed below).

Additionally, we have shown that conversational production and conversational comprehension engage regions outside of the perisylvian language network. The occipital cortex which was engaged during both, possibly as a result of looking at the interlocutor. Producing language in a conversational context compared to baseline (looking at a fixation cross) entailed the recruitment of the superior frontal gyrus (SFG), motor regions, and the medial frontal cortex, while language comprehension in conversation compared to baseline engaged the insula bilaterally. In the more crucial contrast of production and comprehension systems, production recruited regions involved in higher-level sociocognitive processing, such as the bilateral medial prefrontal cortex (mPFC), which has been observed in communicative perspective-taking during utterance planning (Vanlangendonck et al., 2018). Interestingly, no regions outside of the perisylvian language network were more activated for conversational comprehension than conversational production.

Our results extend the functional asymmetry of frontal and temporal regions, previously observed in the controlled production and comprehension experiment of Giglio et al. (2022b), to the crucial conversational setting. Giglio et al. (2022b) used large functional ROIs that extended into regions of the MD network (e.g., the middle frontal gyrus), meaning that the observed asymmetry in their study could have been due to the increased domain-general demands of speaking, as compared to listening. We used data from smaller anatomical ROIs but still observed a similar asymmetry, which, contrary to recent accounts (Hu et al., 2022), supports the notion that frontal and temporal regions in the perisylvian language network are functionally segregated. The observed LIFG specialization for production aligns with the classic model’s characterization of Broca’s area (Geschwind, 1970). However, the increased activation of the LaMTG contradicts the classic model’s strict segregation of comprehension to Wernicke’s area, which is located in the posterior temporal cortex. The observed LIFG specialization is also compatible with the model proposed by Matchin and Hickok (2020), according to which the LIFG supports syntactic processes primarily tied to production, while the LpMTG supports more basic syntactic processing in both production and comprehension. Another plausible explanation that would work as an alternative to the model in Matchin and Hickok (2020) and still attribute system-general syntactic processing to the LIFG (as in e.g. Hagoort, 2016), is that the production of a grammatically adequate utterance requires the speaker to fully build a syntactic representation of that utterance (Garrett, 1988; Bock, 1982), while, in comprehension, listeners may circumvent the need for a full syntactic parse by utilization of lexical information, simple heuristics connecting syntax and semantics (e.g., *the first incoming noun phrase is the agent of an action*), and world knowledge (Ferreira and Patson, 2007; Ferreira et al., 2001; Christianson et al., 2001; Townsend and Bever, 2001; Ferreira, 2003). There might be additional ways in which the production of an in some sense redundant linguistic code is more laborious than comprehension (as there is no need to parse everything in a redundant code). Conceivably, the propensity to employ mechanisms that favor processing speed should be larger during conversational comprehension than during non-interactive comprehension, because of the demands of timing in turn-taking (Levinson and Torreira, 2015; Stivers et al., 2009; Kendrick and Torreira, 2015). Our results point in the direction that these comprehension processes may be supported by the LaMTG/STS.

There is yet another explanation for why we did observe segregation of production and comprehension processes in the LaMTG/STS, while Hu et al. (2022) did not. The anterior temporal cortex has been described as a semantic hub that binds and organizes the semantic features of mental concepts (Hickok, 2009; Lambon Ralph and Patterson, 2008; Patterson et al., 2007). Conversational comprehension does not only require the listener to analyze what is currently being said but also what has previously been said and other contextual information to, e.g., resolve semantic ambiguities, or decode the indirect meaning of an incoming utterance (Grice, 1975). These more complex, semantic processes may not be as strongly recruited in comprehension tasks that do not require the incorporation of previous linguistic information and other contextual cues to infer utterance meaning (e.g., Hu et al., 2022). On the other hand, the whole-brain results in Giglio et al. (2022b) did indicate a stronger activation of the LaMTG in comprehension than production. This observed asymmetry could be ascribed to their use of embedded clauses (e.g. ’The woman saw that the man clapped’), which not only requires more syntactic, but also semantic, combinatorial processes. Moreover, the production task in Hu et al. (2022) was designed to elicit utterances by showing images of events that participants were asked to describe. Conceivably, some aspects of production may even be more demanding (or at least different) in this type of image elicitation, compared to the spontaneous production that takes place during conversation.

We also want to highlight that Matchin and Hickok (2020) describe the posterior, rather than anterior, aspects of the LIFG (the pars triangularis and opercularis) as more crucial to production than comprehension. We have not only found that posterior LIFG activation is stronger in production, but that the anterior LIFG (the pars orbitalis) supports processes more strongly recruited in production than comprehension. The left pars orbitalis clearly plays a key role in semantic retrieval, as it has been shown to activate with increasing semantic conflict at the single-word level (for a review, see Price, 2010), when processing real words vs pseudowords (Nosarti et al., 2010; Schafer and Constable, 2009)), and during object naming vs hearing or reading words (Ekert et al., 2021). A goal for future investigations is to determine whether the production-specific processes subserved by the LIFG pars orbitalis are semantic in nature.

Moving to the extra-linguistic processes of conversation, our results draw attention to the possibility that production, to a greater extent than comprehension, may draw on higher-level sociopragmatic processing. Activation of regions such as the bilateral mPFC, the LMFG, and the bilateral SFG was stronger in production than comprehension. These regions inter alia subserve mechanisms that enable the ability to attribute mental states to oneself and to others – an ability often referred to as theory of mind (Astington et al., 1988; Schurz et al., 2014). The bilateral mPFC and MFG are also part of the MD network, facilitating cognitive control (Camilleri et al., 2018; Diachek et al., 2020; Duncan, 2013; MacGregor et al., 2022; Stiers et al., 2010; Wehbe et al., 2021). In a production planning task (Vanlangendonck et al., 2018), the mPFC and the LSFG were implicated in production planning during communicative vs non-communicative conditions, while the LMFG was implicated in perspective-taking during production planning when the addressee’s visual perspective differed from the speaker’s. In addition, we found that production activation was stronger in the precuneus bilaterally. This activated part of the precuneus (at and around MNI coordinates x: 0 y: −70 z: 55) has been implicated in a task where participants listened to indirect vs direct speech acts (Bendtz et al., 2022).

In comprehension, listeners also rely on social and contextual information to, for example, decode non-literal messages (Bšáková et al., 2015; Bendtz et al., 2022; Grice, 1975). However, it is justifiable to hypothesize that conversational production is socially and pragmatically more demanding than conversational comprehension. Language production, as compared to language comprehension, involves the formulation of an utterance (Levelt, 1993). In formulating an utterance for communicative purposes, the speaker generally strives to meet the goals of communication, which include successfully conveying the intended meaning of the utterance (Arvidsson et al., 2022b; Clark and Murphy, 1982; Papafragou and Grigoroglou, 2019) and avoiding the threatening of one’s own and the other interlocutor’s face (Brown et al., 1987; Goffman, 1967; Schegloff and Sacks, 1973). To meet these goals, speakers need to take into account social and contextual information, such as the specific needs and knowledge states of their interlocutor, while at the same time maintaining relevant linguistic information in memory. Considering this, it is unsurprising that conversational production processes incur a greater cost for regions associated with sociopragmatic and cognitive control processes. While most of these processes may be even stronger e.g. just prior to the onset of production (see

Arvidsson et al., 2022a), it is interesting to note that they also seem to occur during production as such, perhaps as a way of monitoring that the speaker’s intentional goals are met.

Our results provide neurobiological support for the so-called separable view of production and comprehension systems (Kittredge and Dell, 2016), often endorsed in psycholinguistics (Meyer et al., 2016; Gahl and Strand, 2016), indicating that ‘speaking and listening cannot be understood as the same processes running in the opposite directions’ (Meyer et al., 2016). This is, as we have discussed, perhaps most evident for socio-pragmatic processing, but may also apply to utterance/sentence-level syntactic and semantic processes all the way down to the lexicon. At least, the systems appear to differ in the degree to which they rely on common representations. Two limitations of the study relate to core features of conversation. First, although participants of a conversation often take turns talking, it is not uncommon that speech from two individuals overlap. Overlapping speech occurs, e.g., during the transition from one speaker’s turn to another speaker’s turn (Heldner and Edlund, 2010; Sacks et al., 1978). In our study, overlapping speech leads to overlapping production and comprehension events. We do not regard this as a crucial issue, partly because conversational production and comprehension processes naturally overlap, even when there is no overlapping speech (Levinson and Torreira, 2015; Bögels et al., 2018); the listener plans and encodes their upcoming response when the incoming turn from their interlocutor is still unfinished and needs to be monitored. Furthermore, our main result is the asymmetric recruitment pattern of production and comprehension. Such an asymmetry could not be driven but obscured by overlapping production and comprehension processes. Moreover, the very fact that production and comprehension may overlap is a necessary condition for studying these processes in the conversational setting and thus part of our unique contribution.

Second, conversational turns are often short in time (the mean duration in the current study was 1.67 for production and 1.96 for comprehension, see Table 1). Low-duration events (i.e. trials) are not standard in fMRI data analysis, because most MRI studies are based on controlled experiments where the timing of events is predetermined. However, methodological research has shown that responses to stimuli with durations as short as 5 ms can be reliably detected with fMRI (Yeşsilyurt et al., 2008). A larger set of events than the typical 20-50 events per condition (e.g., 48 events/condition in Giglio et al., 2022b) are however needed when studying shorter events. Our study included approximately 3500 events per condition, see Table 1.

In summary, our results extend the asymmetric recruitment pattern of noninteractive production and comprehension processes observed in Giglio et al. (2022b), by showing that conversational production and comprehension diverge with respect to how they modulate the recruitment of regions in the left-lateralized perisylvian language network. The recruitment of LIFG-regions (interestingly including the pars orbitalis) was stronger in production than in comprehension, while the recruitment of the LaMTG/STS was stronger in comprehension than in production. This asymmetric pattern favors descriptions, where (1) frontal regions subserve processes mainly linked to production (Matchin and Hickok, 2020; Matchin et al., 2022) and anterior temporal regions subserve processes that are more strongly recruited in comprehension, or (2) inferior frontal regions support syntactic combinatorial processes that can be circumvented in comprehension (Giglio et al., 2022b). This circumvention may be enabled by the engagement of semantic processes in LaMTG/STS. Finally, the results also indicate that conversational production, even to a greater extent than conversational comprehension, may draw on higher-level sociopragmatic processes, subserved by cortical regions extending outside of the perisylvian language network (possibly to monitor that the speaker’s intentional goals are met).

We have addressed the long-lived yet ongoing debate of whether there is a functional segregation of production and comprehension processes in frontal vs temporal perisylvian language regions, finally including the conversational context. In conclusion, while providing evidence for a functional asymmetry of the two systems, in the expected direction following the Broca-Wernicke model, the temporal functional anatomy suggested by our results do depart from the classic model. This is an example of how investigations of *conversational* production and comprehension will alter models built on isolation paradigms. These results, together with the results on system asymmetries outside the two classical language regions, suggest there may well be differences in knowledge representations across systems, e.g. for socio-pragmatic processes, while common knowledge representations might be used to different extents at the lexico-syntactic level.

## Acknowledgments

This work was supported by the Digital Futures project ’Using Neuroimaging Data for Exploring Conversational Engagement in Human-Robot Interaction’, Riksbankens Jubileumsfond (http://dx.doi.org/10.13039/501100004472 to JU), and the Swedish Collegium of Advanced Studies (to JU).

## References

Arvidsson C, Ekaterina T, Andé P, Uddén J. 2022a. The brain in conversation: Mapping turn-taking, production and comprehension with fmri. In: The Fourteenth Annual Meeting of the Society for the Neurobiology of Language, Philadelphia, USA, October 6-8, 2022.

Arvidsson C, Pagmar D, Uddén J. 2022b. When did you stop speaking to yourself? adolescent development of world knowledge-based audience design. doi:10.31234/osf.io/mrqgv.

Astington JW, Harris PL, Olson DR. 1988. Developing theories of mind. Cambridge: Cambridge University Press.

Austin J. 1973. Speech acts. The Edinburgh Course in Applied Linguistics. 1:37–53.

Bšáková J, van Berkum J, Weber K, Hagoort P. 2015. A job interview in the mri scanner: How does indirectness affect addressees and overhearers? Neuropsychologia. 76:79–91.

Bendtz K, Ericsson S, Schneider J, Borg J, Bšáková J, Uddén J. 2022. Individual differences in indirect speech act processing found outside the language network. Neurobiology of Language. 3:287–317.

Bock JK. 1982. Toward a cognitive psychology of syntax: Information processing contributions to sentence formulation. Psychological review. 89:1.

Bögels S, Casillas M, Levinson SC. 2018. Planning versus comprehension in turn-taking: Fast responders show reduced anticipatory processing of the question. Neuropsychologia. 109:295–310.

Bookheimer S. 2002. Functional mri of language: new approaches to understanding the cortical organization of semantic processing. Annual review of neuroscience. 25:151–188.

Brown P, Levinson SC, Levinson SC. 1987. Politeness: Some universals in language usage. volume 4. Cambridge university press.

Camilleri JA, Müller VI, Fox P, Laird AR, Hoffstaedter F, Kalenscher T, Eickhoff SB. 2018. Definition and characterization of an extended multiple-demand network. NeuroImage. 165:138–147.

Caplan D. 1987. Neurolinguistics and linguistic aphasiology: An introduction. Cambridge University Press.

Caramazza A, Zurif EB. 1976. Dissociation of algorithmic and heuristic processes in language comprehension: Evidence from aphasia. Brain and Language. 3:572–582.

Chomsky N. 2014. Aspects of the Theory of Syntax. volume 11. MIT press.

Christianson K, Hollingworth A, Halliwell JF, Ferreira F. 2001. Thematic roles assigned along the garden path linger. Cognitive psychology. 42:368–407.

Clark HH, Murphy GL. 1982. Audience design in meaning and reference. In: JF Le Ny, W Kintsch, editors. Advances in Psychology. volume9 of Language and Comprehension. North-Holland. p. 287–299. doi:https://10.1016/S0166-4115(09)60059-5.

Diachek E, Blank I, Siegelman M, Affourtit J, Fedorenko E. 2020. The domain-general multiple demand (md) network does not support core aspects of language comprehension: a large-scale fmri investigation. Journal of Neuroscience. 40:4536– 4550.

Duncan J. 2013. The structure of cognition: attentional episodes in mind and brain. Neuron. 80:35–50.

Ekert JO, Lorca-Puls DL, Gajardo-Vidal A, Crinion JT, Hope TM, Green DW, Price CJ. 2021. A functional dissociation of the left frontal regions that contribute to single word production tasks. Neuroimage. 245:118734.

Fedorenko E, Behr MK, Kanwisher N. 2011. Functional specificity for high-level linguistic processing in the human brain. Proceedings of the National Academy of Sciences. 108:16428–16433.

Ferreira F. 2003. The misinterpretation of noncanonical sentences. Cognitive psychology. 47:164–203.

Ferreira F, Christianson K, Hollingworth A. 2001. Misinterpretations of garden-path sentences: Implications for models of sentence processing and reanalysis. Journal of Psycholinguistic Research. 30:3–20.

Ferreira F, Patson ND. 2007. The ‘good enough’ approach to language comprehension. Language and Linguistics Compass. 1:71–83.

Fridriksson J, Fillmore P, Guo D, Rorden C. 2015. Chronic broca’s aphasia is caused by damage to broca’s and wernicke’s areas. Cerebral Cortex. 25:4689–4696.

Gahl S, Strand JF. 2016. Many neighborhoods: Phonological and perceptual neighborhood density in lexical production and perception. Journal of Memory and Language. 89:162–178.

Garrett MF. 1988. Processes in language production. volume 3.

Geschwind N. 1970. The organization of language and the brain: Language disorders after brain damage help in elucidating the neural basis of verbal behavior. Science. 170:940–944.

Giglio L, Ostarek M, Sharoh D, Hagoort P. 2022a. Diverging neural dynamics for syntactic structure building in naturalistic speaking and listening. bioRxiv. :2022– 10.

Giglio L, Ostarek M, Weber K, Hagoort P. 2022b. Commonalities and asymmetries in the neurobiological infrastructure for language production and comprehension. Cerebral Cortex. 32:1405–1418.

Goffman E. 1967. On face-work. Interaction ritual. :5–45.

Grice HP. 1975. Logic and conversation. In: P Cole, JL Morgan, editors. Speech acts. Brill. p. 41–58.

Hagoort P. 2016. Muc (memory, unification, control): A model on the neurobiology of language beyond single word processing. In: Neurobiology of Language. Elsevier. p. 339–347. doi:https://doi.org/10.1016/B978-0-12-407794-2.00028-6.

Heldner M, Edlund J. 2010. Pauses, gaps and overlaps in conversations. Journal of Phonetics. 38:555–568.

Hickok G. 2009. The functional neuroanatomy of language. Physics of life reviews. 6:121–143.

Hickok G. 2022. The dual stream model of speech and language processing. Handbook of Clinical Neurology. 185:57–69.

Hogenhuis A, Hortensius R. 2022. Domain-specific and domain-general neural network engagement during human–robot interactions. European Journal of Neuroscience. 56:5902–5916.

Hu J, Small H, Kean H, Takahashi A, Zekelman L, Kleinman D, Ryan E, Nieto-Castañón A, Ferreira V, Fedorenko E. 2022. Precision fmri reveals that the language-selective network supports both phrase-structure building and lexical access during language production. Cerebral Cortex. 1:21.

Kendrick KH, Torreira F. 2015. The timing and construction of preference: A quantitative study. Discourse Processes. 52:255–289.

Kittredge AK, Dell GS. 2016. Learning to speak by listening: Transfer of phonotactics from perception to production. Journal of Memory and Language. 89:8–22.

Lambon Ralph MA, Patterson K. 2008. Generalization and differentiation in semantic memory: insights from semantic dementia. Annals of the New York Academy of Sciences. 1124:61–76.

Levelt WJ. 1993. Speaking: From intention to articulation. MIT press.

Levinson SC, Torreira F. 2015. Timing in turn-taking and its implications for processing models of language. Frontiers in Psychology. 6:731.

MacGregor LJ, Gilbert RA, Balewski Z, Mitchell DJ, Erzinclioglu SW, Rodd JM, Duncan J, Fedorenko E, Davis MH. 2022. Causal contributions of the domain-general (multiple demand) and the language-selective brain networks to perceptual and semantic challenges in speech comprehension. bioRxiv.

Malik-Moraleda S, Ayyash D, Gallée J, Affourtit J, Hoffmann M, Mineroff Z, Jouravlev O, Fedorenko E. 2022. An investigation across 45 languages and 12 language families reveals a universal language network. Nature Neuroscience. 25:1014–1019.

Matchin W, Basilakos A, Den Ouden DB, Stark BC, Hickok G, Fridriksson J. 2022. Functional differentiation in the language network revealed by lesion-symptom mapping. NeuroImage. 247:118778.

Matchin W, Hickok G. 2020. The cortical organization of syntax. Cerebral Cortex. 30:1481–1498.

Meyer AS, Huettig F, Levelt WJ. 2016. Same, different, or closely related: What is the relationship between language production and comprehension? Journal of Memory and Language. 89:1–7.

Miller EK, Cohen JD. 2001. An integrative theory of prefrontal cortex function. Annual Review of Neuroscience. 24:167–202.

Mollica F, Siegelman M, Diachek E, Piantadosi ST, Mineroff Z, Futrell R, Kean H, Qian P, Fedorenko E. 2020. Composition is the core driver of the language-selective network. Neurobiology of Language. 1:104–134.

Nosarti C, Mechelli A, Green DW, Price CJ. 2010. The impact of second language learning on semantic and nonsemantic first language reading. Cerebral Cortex. 20:315–327.

Pallier C, Devauchelle AD, Dehaene S. 2011. Cortical representation of the constituent structure of sentences. Proceedings of the National Academy of Sciences. 108:2522– 2527.

Papafragou A, Grigoroglou M. 2019. The role of conceptualization during language production: evidence from event encoding. Language, Cognition and Neuroscience. 34:1117–1128.

Patterson K, Nestor PJ, Rogers TT. 2007. Where do you know what you know? the representation of semantic knowledge in the human brain. Nature reviews neuroscience. 8:976–987.

Pickering MJ, Garrod S. 2004. Toward a mechanistic psychology of dialogue. Behavioral and Brain Sciences. 27:169–190.

Pickering MJ, Garrod S. 2007. Do people use language production to make predictions during comprehension? Trends in cognitive sciences. 11:105–110.

Price CJ. 2010. The anatomy of language: a review of 100 fmri studies published in 2009. Annals of the New York Academy of Sciences. 1191:62–88.

R Core Team. 2021. R: A Language and Environment for Statistical Computing. R Foundation for Statistical Computing. Vienna, Austria.

Rauchbauer B, Hmamouche Y, Bigi B, Prevot L, Ochs M, Thierry C. 2020. Multimodal Corpus of Bidirectional Conversation of Human-human and Human-robot Interaction during fMRI Scanning. In: Proceedings of The 12th Language Resources and Evaluation Conference. Marseille, France: European Language Resources Association. p. 661–668.

Rauchbauer B, Nazarian B, Bourhis M, Ochs M, Pévot L, Chaminade T. 2019. Brain activity during reciprocal social interaction investigated using conversational robots as control condition. Philosophical Transactions of the Royal Society B. 374:20180033.

Rolls ET, Huang CC, Lin CP, Feng J, Joliot M. 2020. Automated anatomical labelling atlas 3. NeuroImage. 206:116189.

Rutten GJ. 2022. Broca-wernicke theories: A historical perspective. Handbook of Clinical Neurology. 185:25–34.

Sacks H, Schegloff EA, Jefferson G. 1978. A simplest systematics for the organization of turn taking for conversation. In: Studies in the Organization of Conversational Interaction. Elsevier. p. 7–55. doi:https://doi.org/10.1016/B978-0-12-623550-0.50008-2.

Schafer RJ, Constable T. 2009. Modulation of functional connectivity with the syntactic and semantic demands of a noun phrase formation task: a possible role for the default network. Neuroimage. 46:882–890.

Schegloff EA, Sacks H. 1973. Opening up closings. Semiotica. 8:289–327.

Schurz M, Radua J, Aichhorn M, Richlan F, Perner J. 2014. Fractionating theory of mind: a meta-analysis of functional brain imaging studies. Neuroscience & Biobehavioral Reviews. 42:9–34.

Segaert K, Menenti L, Weber K, Petersson KM, Hagoort P. 2012. Shared syntax in language production and language comprehension—an fmri study. Cerebral Cortex. 22:1662–1670.

Singmann H, Bolker B, Westfall J, Aust F, Ben-Shachar M. 2018. afex: Analysis of factorial experiments [r package]. Retrieved fro m https://cran.r-project.org/package=afex.

Stiers P, Mennes M, Sunaert S. 2010. Distributed task coding throughout the multiple demand network of the human frontal–insular cortex. Neuroimage. 52:252–262.

Stivers T, Enfield NJ, Brown P, Englert C, Hayashi M, Heinemann T, Hoymann G, Rossano F, De Ruiter JP, Yoon KE, Levinson SC. 2009. Universals and cultural variation in turn-taking in conversation. Proceedings of the National Academy of Sciences. 106:10587–10592.

Stockert A, Hormig-Rauber S, Wawrzyniak M, Klingbeil J, Schneider HR, Pirlich M, Schob S, Hoffmann KT, Saur D. 2023. Involvement of thalamocortical networks in patients with poststroke thalamic aphasia. Neurology. 100:e485–e496.

Townsend DJ, Bever TG. 2001. Sentence comprehension: The integration of habits and rules. Cambridge: MIT Press. doi:https://doi.org/10.7551/mitpress/6184.001.0001.

Tremblay P, Dick AS. 2016. Broca and wernicke are dead, or moving past the classic model of language neurobiology. Brain and language. 162:60–71.

Tyler LK, Shafto MA, Randall B, Wright P, Marslen-Wilson WD, Stamatakis EA. 2010. Preserving syntactic processing across the adult life span: the modulation of the frontotemporal language system in the context of age-related atrophy. Cerebral Cortex. 20:352–364.

Vanlangendonck F, Willems RM, Hagoort P. 2018. Taking common ground into account: Specifying the role of the mentalizing network in communicative language production. PloS One. 13:e0202943.

Vazquez A, Bates D, Rosa G, Gianola D, Weigel K. 2010. an r package for fitting generalized linear mixed models in animal breeding. Journal of animal science. 88:497–504.

Vlooswijk M, Jansen J, Majoie H, Hofman P, de Krom M, Aldenkamp A, Backes W. 2010. Functional connectivity and language impairment in cryptogenic localization-related epilepsy. Neurology. 75:395–402.

Wehbe L, Blank IA, Shain C, Futrell R, Levy R, von der Malsburg T, Smith N, Gibson E, Fedorenko E. 2021. Incremental language comprehension difficulty predicts activity in the language network but not the multiple demand network. Cerebral Cortex. 31:4006–4023.

Wernicke C. 1885/1977. Recent works on aphasia. In: GH Eggert, editor. Wernicke’s works on aphasia. The Hague: De Gruyter Mouton. p. 173–205.

Wilson D, Sperber D. 2002. Relevance theory.

Yeşilyurt B, Uğurbil K, Uludğ K. 2008. Dynamics and nonlinearities of the bold response at very short stimulus durations. Magnetic resonance imaging. 26:853–862.

